# T13F2.2 of *C. elegans* is Critical for Chromatin Organization, Autophagy, and Longevity

**DOI:** 10.1101/2025.09.22.677762

**Authors:** Rohil Hameed, Sweta Sikder, Aayushi Agrawal, Kottapalli Srividya, Kalyan Mitra, Madavan Vasudevan, Parijat Senapati, Aamir Nazir, Tapas K. Kundu

## Abstract

Intricately modulated by a spectrum of proteins, Chromatin structure governs gene expression and cellular homeostasis. In *Caenorhabditis elegans*, critical components like the TATA-binding protein *tbp-1* play pivotal roles in orchestrating chromatin dynamics. While the function of many of the interacting partners of tbp-1 is well-understood, our study brings into focus a lesser-known entity, T13F2.2, an unexplored tbp-1 interacting protein and a putative RNA polymerase II transcriptional coactivator. Employing reverse genetics, we found that RNAi-induced depletion of T13F2.2 resulted in pronounced disruptions to nuclear architecture, evidenced by nuclear staining and transmission electron microscopy. Accompanying these structural anomalies, we observed increased autophagy, pointing to cellular stress and a hyperacetylation of the core histones, suggesting potential chromatin decompaction. Notably, multifaceted functional alterations, upon the partial knockdown of the T13F2.2, culminated in a substantial reduction in the worm’s lifespan. Intriguingly, interventions such as administering ROS scavengers and autophagy modulators offered a reprieve from this life-shortening effect. Transcriptomic analysis upon T13F2.2 knockdown revealed upregulation of genes related to autophagy and chromatin remodelling, alongside downregulation of genes involved in longevity pathways and oxidative stress response. This study, thus, not only puts forward the functional implication of an uncharacterized gene in *C. elegans* biology, but also further emphasizes the role of chromatin organization in aging at the organismal level.

## Introduction

The nematode *Caenorhabditis elegans* provides a biologically relevant platform for exploring cellular, molecular, and various functional aspects of organismal biology, particularly in the fields of developmental genetics and neurobiology. The simplicity of this worm, both in its genetic makeup and its physiological processes, provides an ideal framework for investigating complex biological phenomena such as aging, neurodegeneration, and cellular homeostasis (Anon 1986; Kaletta & Hengartner 2006). With its fully sequenced genome and the availability of advanced genetic tools, *C. elegans* offers the opportunity to dissect the roles of specific genes within living organisms. Amongst multiple key processes, chromatin organization, a fundamental aspect of genomic regulation, orchestrates a broad array of biological functions across all organisms (Rankin 2002). In eukaryotes, the dynamic structure of chromatin is a crucial determinant of gene expression, whereby the modification of histone proteins and the remodeling of chromatin architecture facilitate or hinder the access of transcriptional machinery to DNA. The chromatin landscape in *C. elegans* mirrors that of more complex systems, such as humans, making it an apt model for studying epigenetic regulation (Antoshechkin & Sternberg 2007). *C elegans* possesses a highly evolved beads on a string chromatin organization, comprising four core histones and linker histones. Besides core histones and linker histone H1, like many other higher eukaryotic organisms, *C. elegans* chromatin is also functionalized by nonhistone proteins HPL2 (human HP1alpha homologue), and *C. elegans* possesses highly developed epigenetic systems, which include chromatin remodeling enzymes, chromatin modification enzymes, and several noncoding RNAs (Couteau et al. 2002).

In the eukaryotic genome organization, transcription factors also function as bookmarking proteins. One such protein is the transcription initiation factor, TATA-binding protein (TBP). The TBP and associated factors form the TFIID complex, essential for the initiation of transcription by RNA polymerase II (Araya et al. 2014). The TBP is also essential for RNA polymerase III and RNA polymerase I-mediated transcription initiation.

In this study, we have tried to understand the functional significance of an uncharacterized *C. elegans* protein T13F2.2, which putatively (source: Wormbase-interaction network of tbp-1) interacts with tbp-1, the nematode ortholog of the human TBP (Huang et al. n.d. 2009). We have found that T13F2.2 is critical for the lifespan of the worm. The knockdown of T13F2.2 results in higher autophagy at the organizational level. In the partial absence of it the chromatin organization dramatically alters (open conformation) with concomitant alteration of the epigenetic landscape. The RNA seq data reveal several important pathways that could be regulated by T13F2.2.

## Results

### T13F2.2 Knockdown Reduces Lifespan without Affecting Development or Morphology

To elucidate the physiological role of *T13F2.2*, we carried out RNAi-induced knockdown in the N2 strain of *Caenorhabditis elegans*. The RNAi-mediated reduction of *T13F2.2*, achieving knockdown efficiency between 60-80% across various experiments, did not result in developmental lethality or delays in progression to the L4 larval stage and subsequently to the adult stage. This indicates that *T13F2.2* is not essential for early developmental milestones. Morphological examination of treated worms revealed no abnormalities in key structural features, including the anterior and posterior pharyngeal bulbs and body musculature, confirming that developmental morphology remains intact **(Fig. 1a-b)**. However, while studying the lifespan of worms, it was observed that the impact of T13F2.2 knockdown on lifespan was significant. Using the N2 strain with the AAK-2 mutant strain as a negative control, we observed a marked reduction in the median lifespan from 16 days in control worms to 10 days in worms subjected to *T13F2.2* knockdown, thus leading to a 37.5% decrease in median lifespan **(Fig. 1c-d))**. This substantial reduction underscores the critical role of *T13F2.2* in maintaining the lifespan of *C. elegans*. Further investigation into the gene expression dynamics associated with *T13F2.2* was performed using quantitative real-time PCR.

**Fig. 1:**
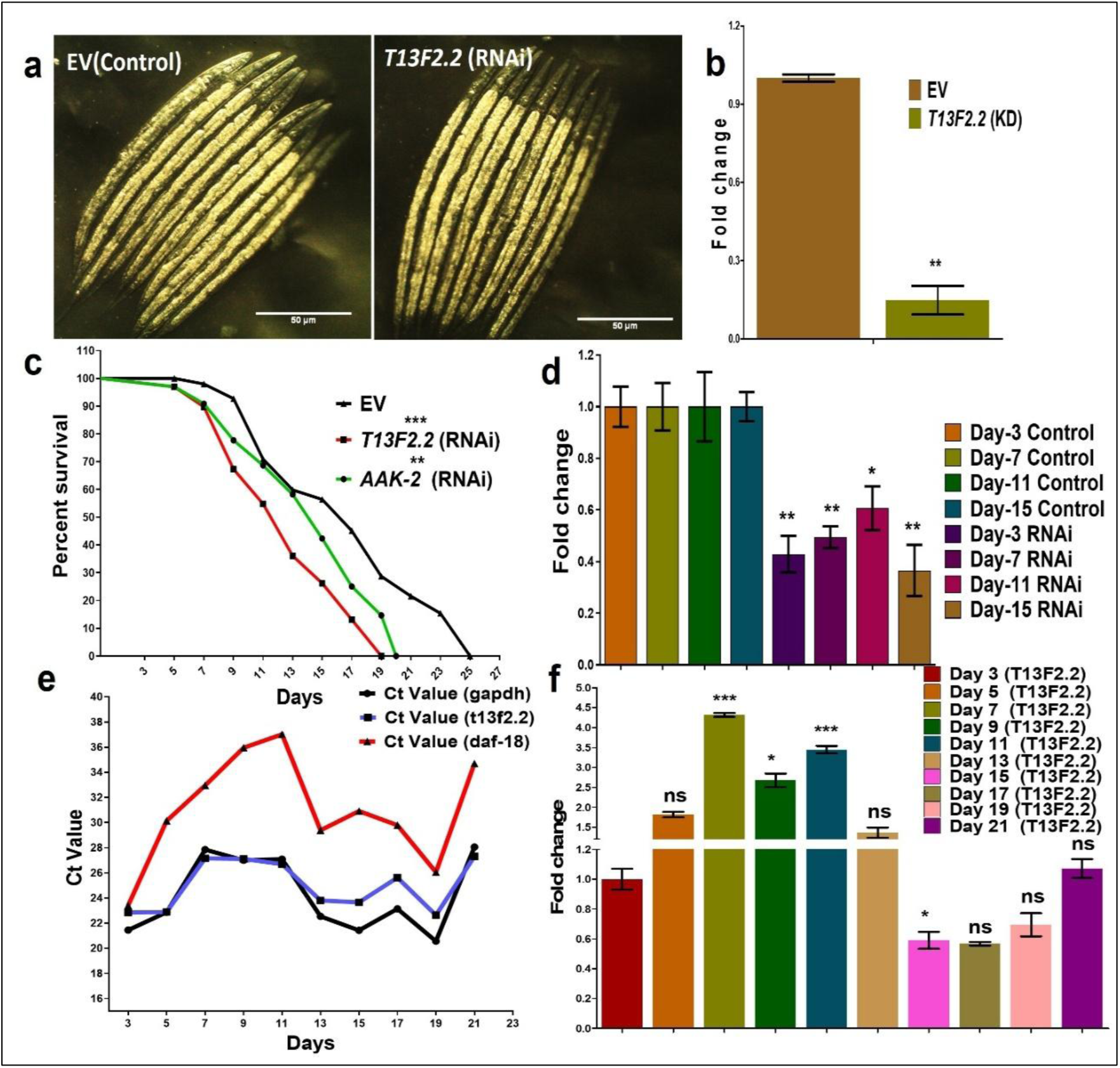
Knockdown of *T13F2.2* is non-lethal, shows no developmental abnormality or delay, but significantly reduces life span in *C. elegans:* **(a)** Wild type (N2 strain) of *C. elegans* showing normal body morphology at young adult stage in, control group (EV) treated with empty vector and after RNAi of *T13F2.2*; all developmental endpoints were unaltered after RNAi. (Magnification-10x), 50um scale bar **(b)** Bar graph depicting a significant decrease in the *T13F2.2* mRNA levels after RNAi of *the T13F2.2 gene,* quantified by q-PCR vs Control. P-Value 0.0028. **(c)** Kaplan-Meier survival curve shows marked decrease in the life span of N2 worms in the knockdown condition of *T13F2.2, AAK-2,* as compared to control conditions (N2 raised on *E. coli* strain EV, RNAi groups fed on HT115) (n=100, P-value, <0.0001) **(d)** Column graph represents RNAi effect at respective days, P-value 0.0023 **(e)** Line graph representing mRNA expression pattern of *T13F2.2*, *GAPDH,* and *Daf-18* quantified via q-PCR across the life span of *C. elegans.* **(f)** Column graph representing change in *T13F2.2* mRNA levels across the life span of *C. elegans* as compared to *T13F2.2* mRNA levels of the young adult worm, P-Value 0.0156.

We studied the expression levels of *T13F2.2*, alongside GAPDH as an internal control and *daf-18*, a gene unrelated to *T13F2.2*’s function, as a functionally neutral gene to provide context for the observed changes. The expression of *T13F2.2* varied significantly throughout the lifespan of the worms, peaking on days 7, 9, and 11 post-hatching and showing a marked decrease by day 15 of the lifespan. Notably, fluctuations in *T13F2.2* expression on days 5, 13, 17, 19, and 21 were not statistically significant **(Fig. 1e-f)**. The mRNA expression pattern across the lifespan is depicted, and changes in T13F2.2 mRNA levels compared to the levels in young adult worms are represented.

### Knockdown of *T13F2.2* leads to induction of autophagy

We further carried out investigations into the physiological impacts of *T13F2.2* knockdown in *Caenorhabditis elegans*, which revealed significant changes in autophagy, a critical cellular process associated with lifespan within the worms, as well as other organisms. To assess these changes, we utilized the DA2123 strain of *C. elegans*, which expresses GFP-tagged autophagosomes in hypodermal seam cells. Photomicrography of these cells showed that *T13F2.2* knockdown via RNAi led to a marked increase in autophagosome formation, as evidenced by intensified fluorescence signaling within the seam cells **(Fig. 2a-b)**. For further validations, we employed the BC12921 strain of the worms, which expresses autophagy marker SQST-1 tagged with green fluorescence protein (GFP in strain SQST-1(P62): GFP. In this strain, RNAi of T13F2.2 resulted in a significantly decreased fluorescence, thus depicting a decrease in SQST-1 levels, which further indicates an upregulation of autophagy. This finding supports the hypothesis that T13F2.2 knockdown enhances autophagy **(Fig. 2e-f)**. To understand the broader implications of T13F2.2 knockdown on autophagy-related gene expression, we analyzed mRNA levels of several key autophagy genes in the N2 strain. Our results showed that upon T13F2.2 knockdown, the mRNA levels of *bec-1, lgg-1, atg-3, atg-4.1,* and *atg-9* significantly increased, while the levels of *unc-51, lgg-2, sqst-1,* and *atg-18* decreased **(Fig.2i)**. This differential expression pattern highlights the complex regulation of autophagy pathways triggered by the loss of T13F2.2.

**Fig. 2:**
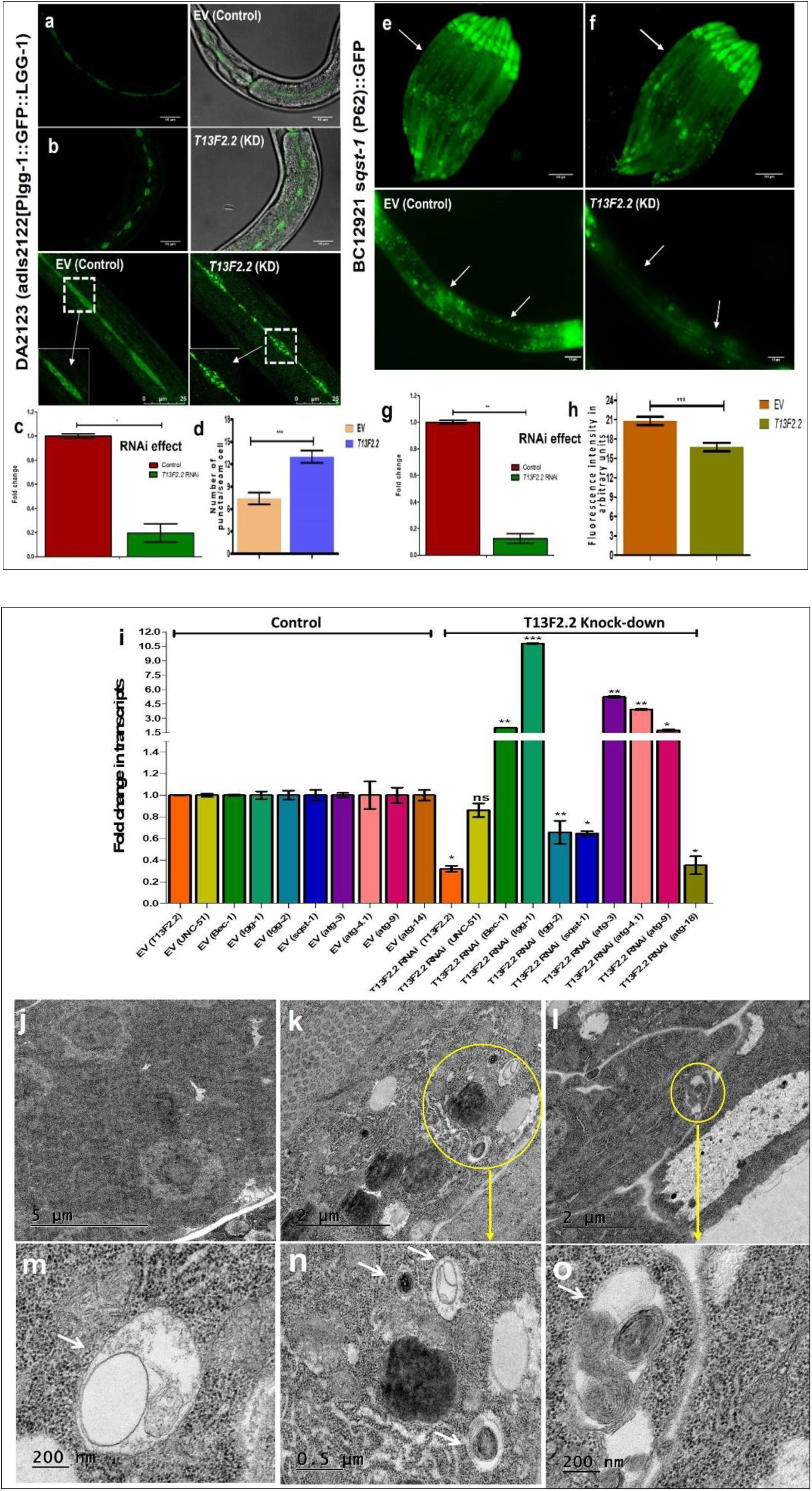
Knockdown of *T13F2.2* leads to induction of autophagy: Representative photomicrographs (a) Control (EV) group, and **(b)** Knockdown of *T13F2.2.* **(c)** the column graph shows the RNAi effect, **(d)** the significant increases in the number of autophagic puncta (having LGG-1::GFP) in seam cells of transgenic *C. elegans* DA2123 (adIs2122[Plgg-1::GFP::LGG-1) (n=10, P-Value, 0.0001)**(e)** Representative images of transgenic *C. elegans* Strain BC12921 (s*qst-1* (P62):: GFP) of Control (EV) group and **(f)** knockdown of *T13F2.2* observed. **(g)** The column graph shows the knockdown effect of *T13F2.2*, **(*h*)** marked decrease in the expression of autophagic receptor SQST-1 (orthologue of human p62). (n=30, P-Value 0.0003) **(i)** Graphical representation of mRNA expression levels for autophagy marker genes (*unc-51, bec-1, lgg-1, lgg-2, sqst-1, atg-3, atg-4.1, atg-9 and atg-18*) quantified by q-PCR employing N2 strain after RNAi of *T13F2.2* with respect to control. (P-Value, 0.0097) **(j-o)** Autophagic structures upon loss of *T13F2.2*; TEM images of *C. elegans* (N2) (EV) control **(j)** group showing normal cellular architecture whereas worms **(k-o)** silenced (RNAi) with *T13F2.2* gene exhibit numerous autophagosomes (white arrows) consisting of double membrane vacuoles with cargo.

Transmission electron microscopy (TEM) provided a further detailed view of the autophagic structures at the cellular level. In worms where T13F2.2 was knocked down, numerous autophagosomes characterized by double-membrane vacuoles were observed, indicating a heightened autophagic response. This contrasted with the control group, which displayed normal cellular architecture with fewer autophagic structures **(Fig. 2j-o)**. These observations collectively demonstrate that the knockdown of T13F2.2 not only reduces lifespan but also significantly induces autophagy in *C. elegans* (**SI.Ia-f)**. The increase in autophagic activity following T13F2.2 knockdown suggests a potential compensatory mechanism that the cells might invoke in response to the reduced functional capacity of other cellular processes. This work provides critical insights into the role of T13F2.2 in regulating autophagy and its implications for cellular health and longevity.

### Knockdown of T13F2.2 Causes Redox Imbalance, Promoting Autophagy and Shortening Organismal Lifespan

Exploring further into the cellular consequences of T13F2.2 knockdown in *Caenorhabditis elegans*, we assessed the impact on the cellular redox state, a critical factor influencing both cell survival and stress responses. Utilizing the H_2_DCFDA method, we measured reactive oxygen species (ROS) levels in the N2 strain post-RNAi treatment. Our findings revealed a significant elevation in ROS levels, indicating a marked redox imbalance induced by T13F2.2 knockdown **(Fig. 3a)**. This increase in ROS was quantified and showed a notable enhancement compared to controls, underlining the stress condition triggered by the gene silencing.

**Fig. 3:**
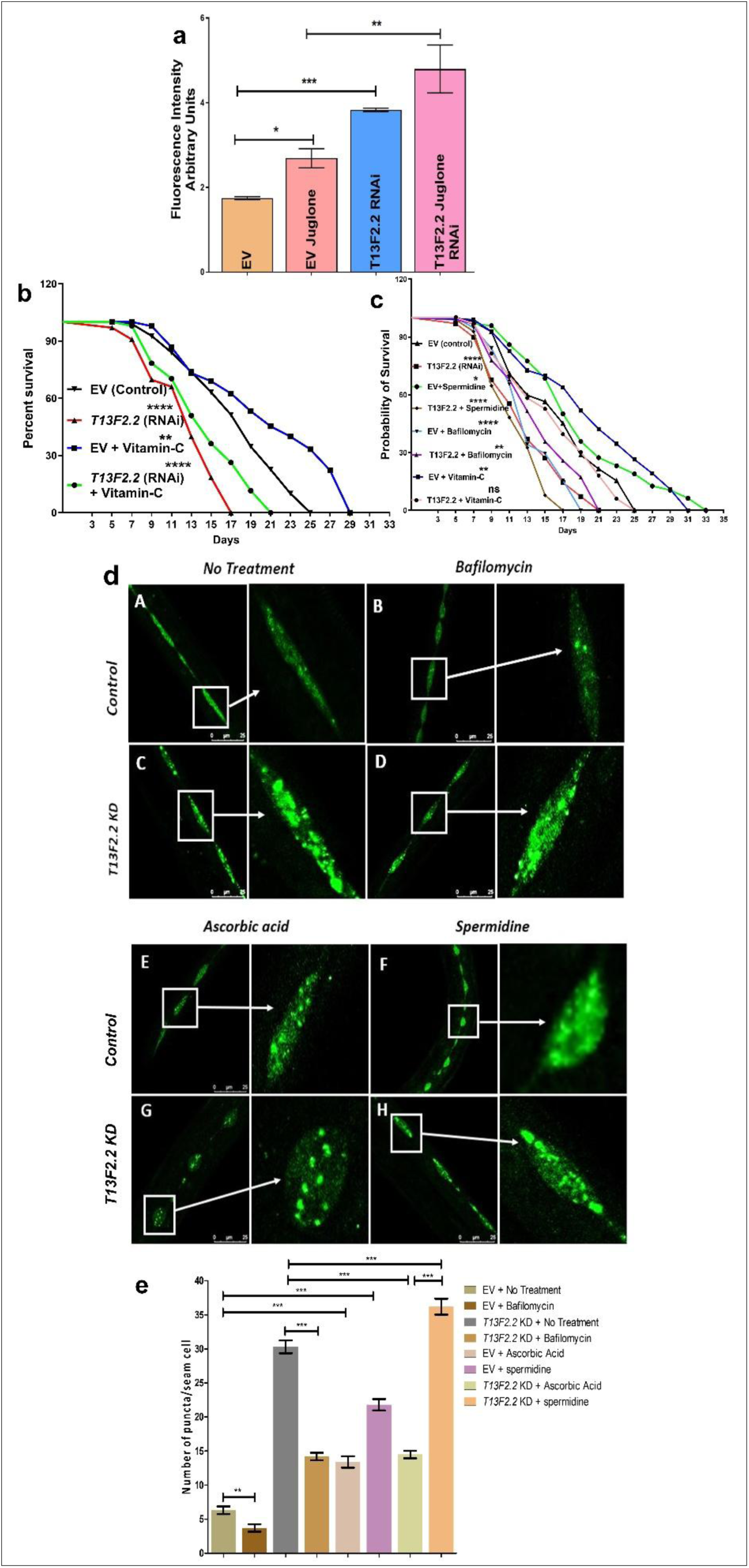
Redox disbalance and induced autophagy leads to reduction in longevity of worms after RNAi of *T13F2.2* **(a)** Significantly elevated ROS levels in N2 worms of *C. elegans* after RNAi of *T13F2.2,* (n=100, P-Value, < 0.0001)**(b)** Kaplan-Meier survival curve shows the restoration of normal life span of wild type in the presence of ascorbic acid after knockdown of *T13F2.2* which was significantly reduced under *T13F2.2* knockdown condition, (n=100, P-Value, < 0.0001) **(c)** The comparative analysis of survival curve shows a significant decrease in the life span of wild type worms in both *T13F2.2* knockdown condition and in the presence of spermidine in the knockdown condition of *T13F2.2*. Worms treated with bafilomycin and ascorbic acid, along with the knockdown of T13F2.2, exhibited a restored life span as compared to their respective controls. **(d)** Restoration of redox balance by ascorbic acid decreases autophagy, which is otherwise increased under RNAi conditions of T13F2.2: Transgenic *C. elegans* DA2123 strain (adIs2122 [Plgg-1:: GFP::LGG-1) exhibits autophagic puncta under **(A)** normal conditions and (**B**) after treatment with Bafilomycin. RNAi of *T13F2.2* worms leads to a significant (**C**) increase in the number and size of autophagic puncta, which is (**D**) decreased in the presence of bafilomycin. (**E**) Treatment of ascorbic acid in the control alters the number of autophagic puncta with respect to the untreated control. (**F**) Spermidine increases its levels further compared to the untreated control. (**G**) Restoration of redox balance by treatment with Ascorbic acid reverses the effect of *T13F2.2* RNAi in terms of an increase in the number of autophagic puncta. (**H**) When worms with knockdown of T13F2.2 are further challenged with autophagy inducer spermidine, the number and size of autophagic puncta are significantly increased. **(e)** The column graph represents the quantification of autophagic puncta in images A-H (n=10, P-Value,0.0010).

Given the established link between redox imbalance and autophagy, we hypothesized that modulating these pathways could mitigate the observed lifespan reduction. Treatment with the antioxidant ascorbic acid effectively countered the lifespan decrease associated with T13F2.2 knockdown. The Kaplan-Meier survival curve demonstrated a restoration of lifespan in ascorbic acid-treated worms to levels comparable to wild-type, underscoring the antioxidant’s protective role **(Fig. 3b)**. Furthermore, we investigated the role of autophagy modulators. The inducer spermidine exacerbated the lifespan reduction when combined with T13F2.2 knockdown, significantly decreasing lifespan beyond the effect of knockdown alone **(Fig. 3c)**. Conversely, treatment with the autophagy inhibitor bafilomycin increased the lifespan, indicating a reversal of the harmful effects induced by T13F2.2 knockdown. The intricate relationship between redox balance and autophagy was further elucidated through our experimental interventions. By assessing autophagic puncta in the DA2123 strain, which displays GFP-tagged autophagosomes, we observed that ascorbic acid treatment not only restored redox balance but also reduced the heightened autophagic activity induced by T13F2.2 knockdown **(Fig. 3d)**. Spermidine treatment, in contrast, led to an increase in both the number and size of autophagic puncta, while bafilomycin treatment decreased these markers **(Fig. 3e)**. Taking them together, these results highlight the intertwined roles of redox imbalance and autophagy in modulating lifespan following T13F2.2 knockdown. The observed redox imbalance could be attributed to disruptions in cellular homeostasis mechanisms, where loss of T13F2.2 potentially impairs the cells’ ability to manage oxidative stress effectively (Schieber & Chandel 2014). This disruption likely leads to an overactivation of autophagy as a compensatory mechanism, which, while initially adaptive, becomes detrimental over time.

### T13F2.2 Localization and Its Impact on Nuclear Architecture in *C. elegans*

T13F2.2, predicted as a coactivator for RNA polymerase II, shares notable functional similarities with the human positive transcriptional coactivator PC4, which is integral to chromatin organization and gene expression regulation (Source: Wormbase/Protein domains). A comparative analysis of the gene and protein sequences between T13F2.2 and human PC4 revealed 54% similarity, suggesting that T13F2.2 may play a similar regulatory role in *C. elegans* **(SI.IIa-d)**. Notably, PC4 is recognized not only for its role in transcription but also as a negative regulator of autophagy, highlighting its critical function in maintaining genomic integrity and cellular homeostasis. To assess the consequences of T13F2.2 knockdown on nuclear architecture, we performed DAPI staining on the N2 strain of *C. elegans*. This approach revealed pronounced distortions in nuclear architecture and potential chromatin decompaction in the absence of T13F2.2. Quantitative analysis showed a 12-fold decrease in DAPI fluorescence intensity in T13F2.2 knockdown worms compared to controls, indicating significant nuclear content reorganization **(Fig. 4a-d)**. Further molecular investigations utilized human PC4 polyclonal antibodies, which identified a protein band corresponding to T13F2.2 in whole cell extracts of *C. elegans* through Western blotting. The intensity of this band was markedly reduced in extracts from T13F2.2 RNAi-treated worms, supporting the antibody’s specificity and the protein’s presence in *C. elegans* **(Fig.SI-III)**. Immunohistochemistry experiments confirmed that T13F2.2 is predominantly localized in the nuclear periphery, specifically in regions of heterochromatin, forming a structure resembling a ring **(Fig. 4h-o)**. This pattern suggests that T13F2.2 may be involved in maintaining the integrity of peripheral heterochromatin and possibly the overall nuclear architecture. Our further experiments with transmission electron microscopy provided detailed insights into the nuclear alterations caused by T13F2.2 deficiency **(Fig. 4p-s)**. Control worms displayed typical nuclear morphology with a regular, spherical nucleus, evenly distributed heterochromatin, a distinct nucleolus, and a well-maintained nuclear envelope. In contrast, worms with T13F2.2 knockdown exhibited a lobulated and malformed nucleus, with noticeable disruptions in the nuclear envelope and a clear loss of peripheral heterochromatin **(Fig.SI-IV)**.

**Fig. 4:**
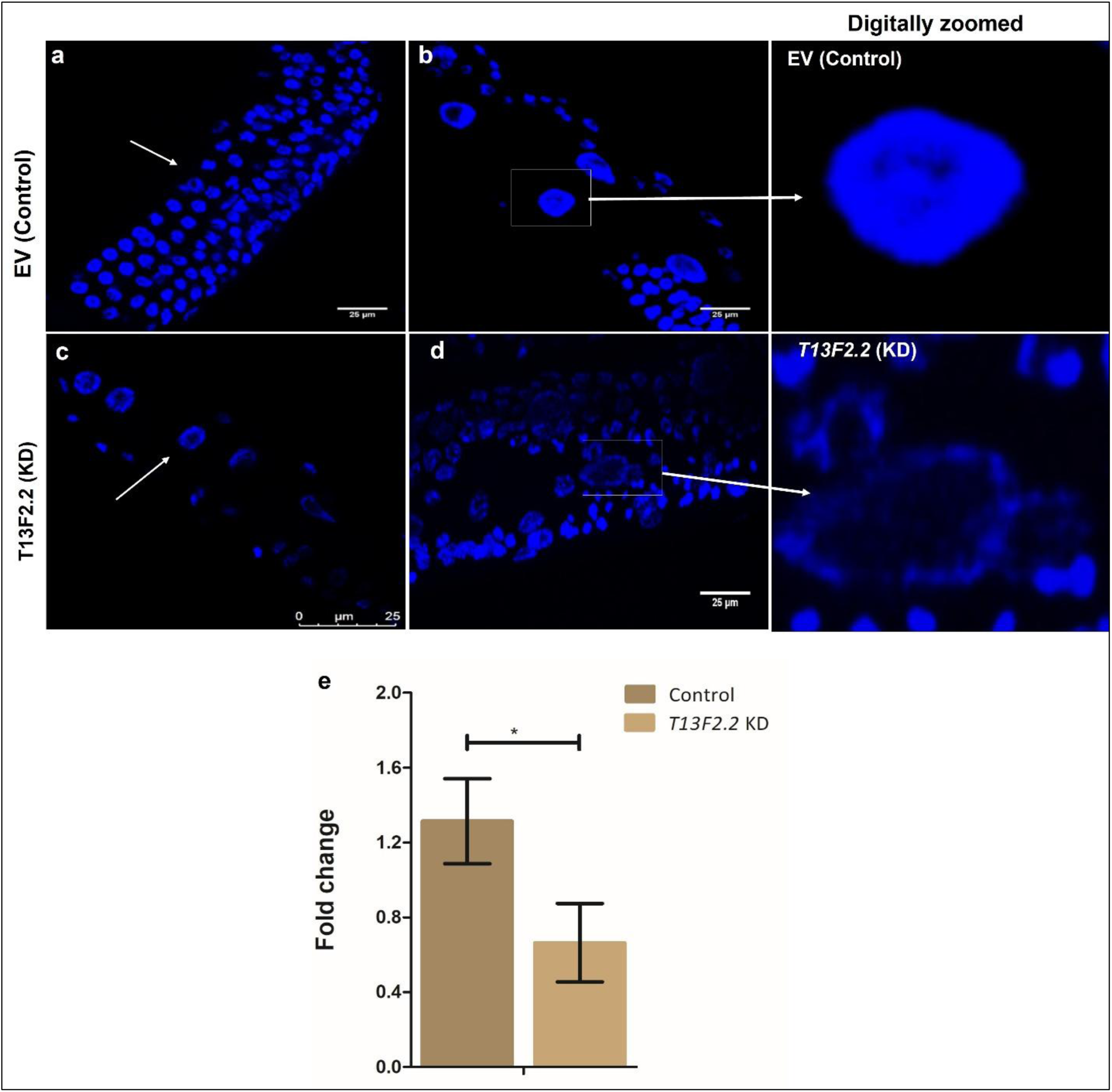
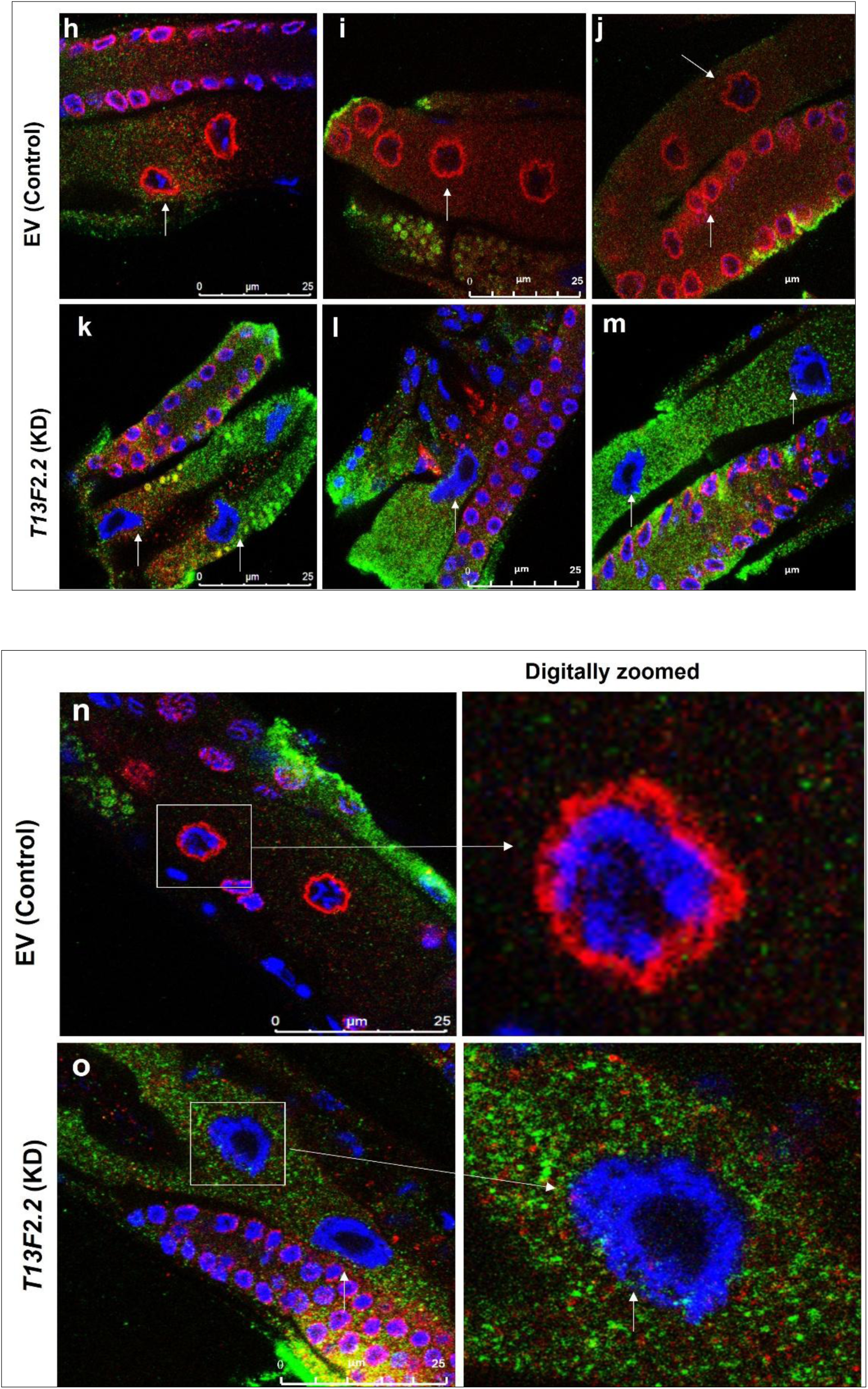
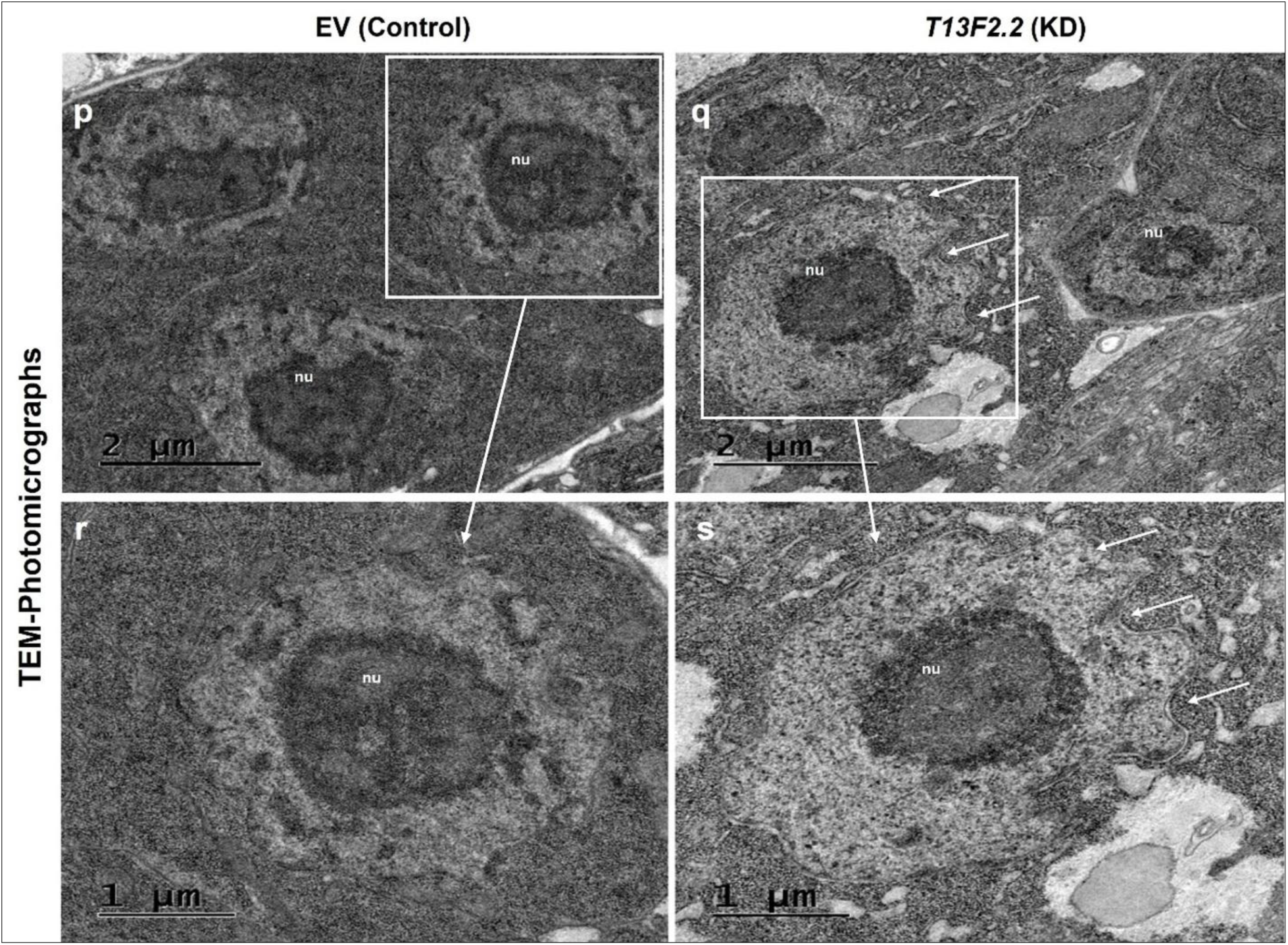
Silencing of *T13F2.2* alters nuclear architecture in *C. elegans*: **(a, b) DAPI-stained images of wild-type (N2) worms, and (c, d)** Decompaction of chromatin upon the knockdown condition of the *T13F2.2* gene. (n=10, P-value, 0.0057) **(h-o)** for Immunohistochemistry we have used the human PC4 antibody which clearly shows, **(h, i, j and n)** the nuclear localization of T13F2.2 protein (red) in Control (EV) and **(k, l, m and o)** destabilized nuclear architecture, pointed as white arrows in the knockdown condition of *T13F2.2* gene. **(p-s):** Transmission electron micrographs of resin-embedded and thin-sectioned *C. elegans* revealing fine structures of nuclei from control and knockdown worms, employing the N2 (wild type) strain. **(r)** Represents photomicrograph acquired at higher magnifications of control worms **(p); (s)** Represents photomicrograph acquired at higher magnifications of knockdown of *T13F2.2* worms **(q).**

These findings collectively demonstrate that T13F2.2 is crucial for maintaining proper nuclear architecture in *C. elegans*. Its absence leads to significant nuclear and chromatin organization disruptions, potentially impairing essential cellular functions and contributing to cellular stress. The resemblance of T13F2.2’s function to that of human PC4 underscores the evolutionary conservation of chromatin organizers and their critical roles in cellular physiology. This study not only highlights the significance of T13F2.2 in nuclear integrity but also provides a foundation for further exploration into its role in transcriptional regulation and cellular stress responses.

### Silencing of T13F2.2 Alters Epigenetic Landscape in *C. elegans:*

We have found that T13F2.2 plays a critical role in maintaining the heterochromatin architecture of the genome in *Caenorhabditis elegans*. To determine its interaction with histones, we employed an antibody pull-down based immunoprecipitation assay with the N2 strain protein lysate. The results revealed that T13F2.2 physically interacts with core histones H2A and H3, as well as the linker histone H1, but not with histones H2B and H4 **(Fig. 5a)**. This selective interaction suggests that T13F2.2 influences genome organization by directly associating with specific components of the nucleosome, particularly those involved in the structural integrity of heterochromatin.

**Fig. 5:**
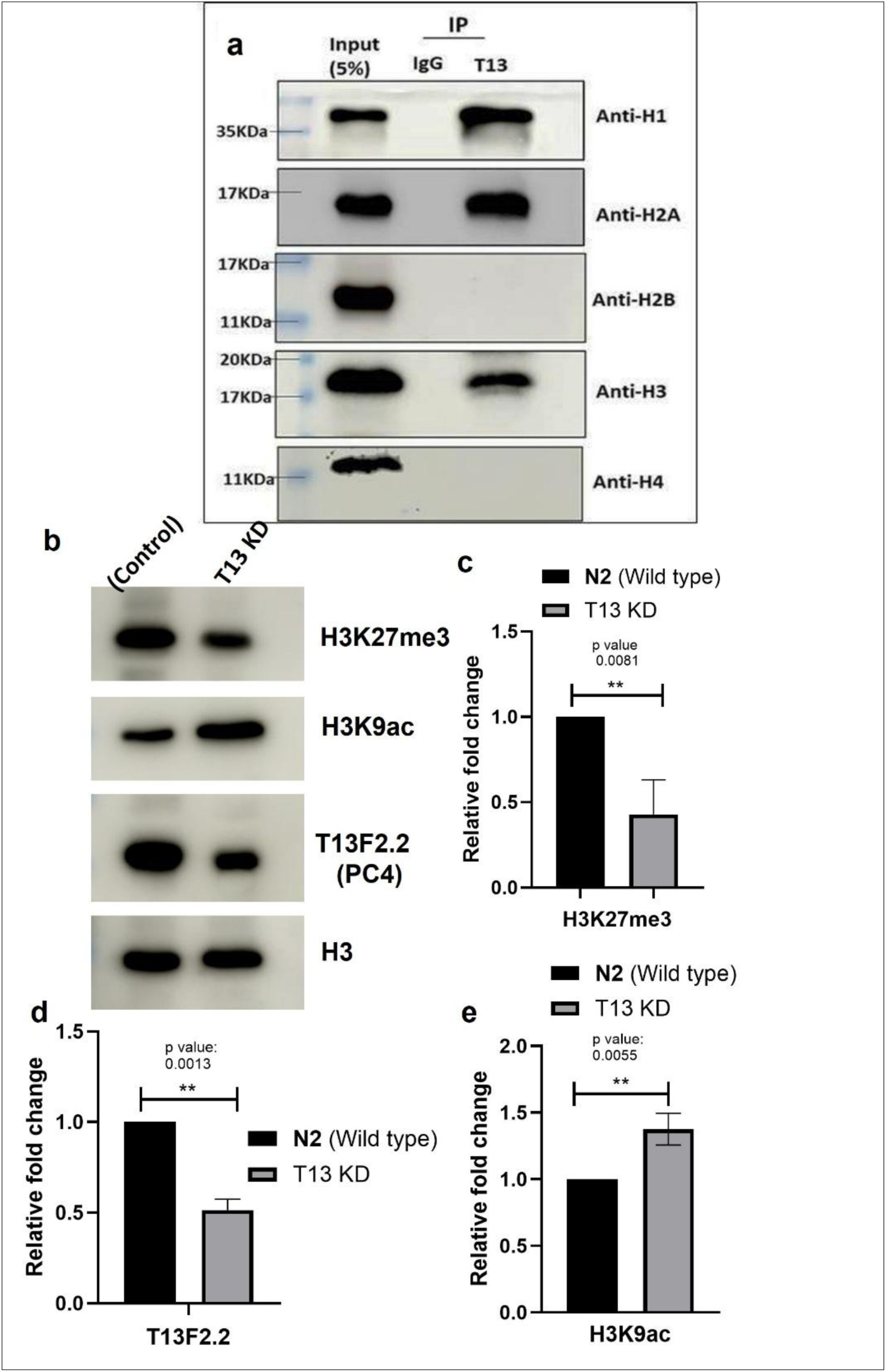
(a) Representative blot of Co-Immunoprecipitation of T13F2.2 using the human PC4 antibody, shows T13F2.2 has physical interaction with H1, H2A, and H3 proteins and not with H2B and H4. **(b-f)** Silencing of *T13F2.2* alters epigenetic landscape: **(b)** Histone modifications marks between control (EV) and *T13F2.2* Knockdown; representative blot and bar graph for its **(c)** significant decrease in H3K27me3, **(d)** densitometry quantification shows a significant decrease in T13F2.2 protein levels and **(f)** significant increase in H3K9ac acetylation levels after RNAi of *T13F2.2* in *C. elegans* N2 strain.

Given the role of T13F2.2 in chromatin compaction, its silencing was hypothesized to alter the epigenetic landscape, potentially affecting gene expression regulation. Immunoblotting analysis conducted to assess levels of known epigenetic acetylation markers associated with transcriptional activation indicated significant changes post-T13F2.2 RNAi. Notably, there was a marked increase in H3K9Ac and a decrease in H3K27me3, upon T13F2.2 knockdown, suggesting a shift towards a more open chromatin state conducive to active transcription **(Fig. 5b)**. Densitometry quantification from Western blots showed a significant decrease in T13F2.2 protein levels in the knockdown samples, correlating with alterations in histone modification patterns. These changes, quantified to show significant shifts in histone acetylation markers, with increased H3K9Ac levels and decreased H3K27me3 levels, support the disruption of chromatin compaction **(Fig. 5d-f)**.

These findings collectively indicate that T13F2.2 is crucial not only for maintaining heterochromatin but also impacts the broader epigenetic landscape of *C. elegans*. The alteration in histone modification patterns following T13F2.2 knockdown underscores its functional importance as a chromatin-associated protein, crucial for both structural integrity and transcriptional regulation within the cell. The disruption of this regulation likely contributes to the phenotypic changes observed upon T13F2.2 silencing, including altered lifespan and increased autophagy, by impacting gene expression through changes in the epigenetic environment.

### Implications of T13F2.2 Loss on Longevity, Autophagy, and Nuclear Organization Genes

The knockdown of T13F2.2 in *C. elegans* resulted in significant alterations in lifespan, autophagic activity, and nuclear architecture, highlighting its role as a critical non-histone nuclear protein. To explore the broader genomic impact of T13F2.2 loss, we conducted whole-worm transcriptomic profiling, which was pivotal for identifying both the genetic actors involved and the pathways affected by T13F2.2 modulation. RNA-seq analysis, using both empty vector (EV) and OP50 as controls, demonstrated high concordance between replicates and significant differences in gene expression between the control and T13F2.2 knockdown (KD) groups, with an 88% variance observed. The analysis revealed a comprehensive alteration in gene expression, with 4112 genes upregulated and 3654 genes downregulated when using EV as a control. With OP50 as the control, slightly fewer genes were differentially regulated. Overall, the comparison between both control conditions showed that 2855 genes were consistently upregulated, and 2537 genes were consistently downregulated across the conditions **(Fig. 6a)**. Pathway analysis on the upregulated genes highlighted significant enrichment in biological pathways such as Chromatin remodeling, Chromatin organization, and Autophagy **(Fig. 6b-c)**. Notably, genes involved in autophagy, including the *vps-39* gene, which is crucial for vesicle-mediated protein trafficking to lysosomes, were found to be upregulated. *Vps-39* is also a part of the HOPS endosomal tethering complex, which plays roles in autophagic processes **(Fig. 6d)**. Conversely, pathway analysis of the downregulated genes indicated a substantial decrease in several metabolic pathways, particularly those involving enzymes with oxidoreductase activity and glutathione transferase activity. This includes the significant downregulation of *gst-2*, a glutathione S-transferase gene crucial for detoxification processes by conjugating reduced glutathione to electrophilic substrates, suggesting a compromised detoxification capability in the worms. Other critical enzymes like superoxide dismutases (*sod-1*, *sod-2*), catalases (*ctl-1*, *ctl-2*), and proteins involved in oxidative stress management, such as Peroxiredoxin and thioredoxin (prdx-2, prdx-6), also showed reduced expression, pointing to an increased vulnerability to oxidative stress **(Fig. 6e-f)**. These transcriptomic findings are consistent with the observed phenotypic changes post-T13F2.2 knockdown, notably the decreased longevity through elevated reactive oxygen species levels and disrupted cellular homeostasis **(Fig. 6g)**. These data collectively underscore the significant role T13F2.2 plays in promoting longevity by maintaining crucial cellular processes such as autophagy, chromatin organization, and oxidative stress management. The detailed heatmap, volcano plot, and genome browser tracks provided further illustrate the depth and breadth of the genetic and epigenetic alterations induced by T13F2.2 loss, providing a comprehensive view of its impact on *C. elegans* physiology and health.

**Fig 6:**
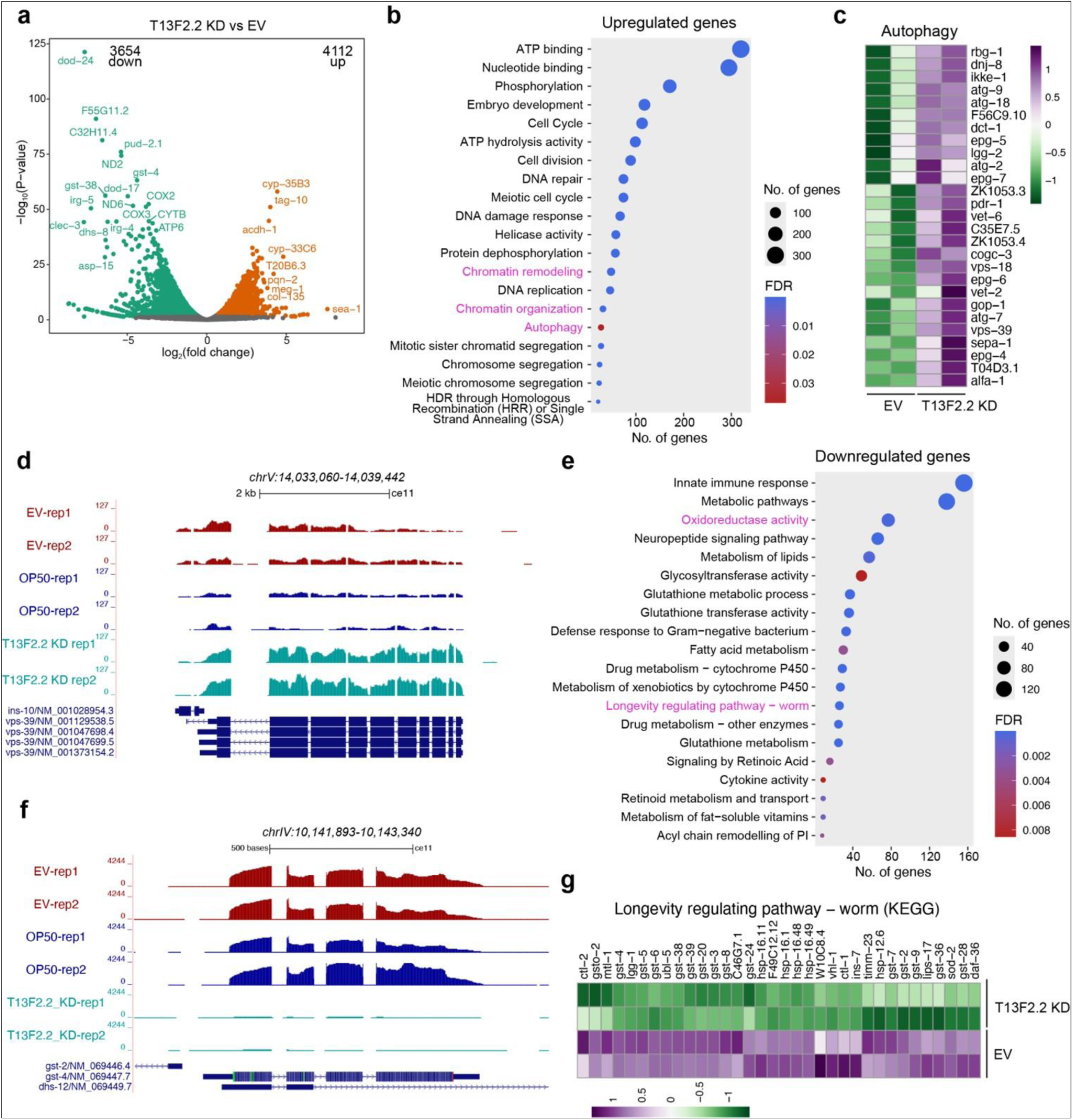
(a) Volcano plot showing the differentially expressed genes between T13F2.2 KD and EV groups. Upregulated genes are shown in orange, and downregulated genes are shown in green. **(b)** Pathways enriched among upregulated genes. The size of the dot indicates the number of genes upregulated, and the color indicates the FDR value. **(c)** The heatmap shows genes involved in Autophagy and the expression across the EV and T13F2.2 KD samples. Z score measures the standard deviation from the mean of log-transformed counts for each gene across all samples. **(d)** Genome Browser tracks show the vps-39 gene locus and the RNA-seq signals across the indicated samples. **(e)** Pathways enriched among downregulated genes. The size of the dot indicates the number of genes downregulated, and the color indicates the FDR value. **(f)** Genome Browser tracks show the gst-2 gene locus and the RNA-seq signals across the indicated samples. **(g)** Heatmap shows genes involved in the Longevity regulating pathway in worms from the KEGG database and the expression across the EV and T13F2.2 KD samples. Z score measures the standard deviation from the mean of log-transformed counts for each gene across all samples.

## Discussion

Aging is a multifactorial process influenced by genetic, epigenetic, and environmental cues that converge on key regulators of cellular homeostasis. Chromatin organization and its interplay with stress response pathways have emerged as critical determinants of longevity and health span (Debès et al. 2023) (Silva-García et al. 2023). In this study, we identified and characterized T13F2.2 as a previously unannotated but functionally important nuclear protein in *Caenorhabditis elegans*, influencing multiple cellular and organismal processes. The depletion of T13F2.2 not only perturbed nuclear morphology and chromatin organization but also compromised longevity, indicating its non-redundant and essential role in organismal homeostasis and related critical processes. In essence, our findings point towards an evolutionarily conserved function of this protein family in chromatin organization, transcriptional control, and cellular stress modulation. The predicted physical interaction of T13F2.2 with *tbp-1*, the *C. elegans* homolog of human TATA-binding protein (TBP), also suggests its role as a transcriptional co-regulator (Talbert & Henikoff 2010). These insights expand the spectrum of chromatin-associated proteins known to shape the aging trajectory of multicellular organisms.

A key highlight of our findings is the robust induction of autophagy upon T13F2.2 knockdown, a process intricately linked with stress responses and longevity. *C. elegans* has long served as a robust and amenable model to decipher lifespan-regulatory mechanisms, including insulin/IGF-1 signaling, dietary restriction, mitochondrial stress, and autophagy (Wilhelm et al. 2017). While the phenomenon of autophagy is generally considered protective, excessive or dysregulated autophagic flux - as observed in the worms with knockdown of T13F2.2 via RNAi - can be deleterious, leading to cellular exhaustion and reduced longevity (Papai et al. 2011).

Interestingly, we observed upregulation of several autophagy genes (e.g., *bec-1*, *lgg-1*, *atg-3*, *atg-4.1*, *atg-9*) and downregulation of others (e.g., *sqst-1*, *unc-51*, *atg-18*), suggesting a non-canonical and imbalanced autophagy program upon knockdown of T13F2.2. Concurrently, a marked increase in ROS levels points to redox imbalance as a potential upstream trigger. The ability of ascorbic acid (an antioxidant) to restore lifespan and suppress autophagic activity reinforces this linkage. In contrast, autophagy induced by spermidine worsened phenotypes, while bafilomycin offered partial rescue, confirming the causative role of excessive autophagy in T13F2.2-deficient animals. These data position T13F2.2 as a novel aging regulator integrating chromatin state, oxidative stress, and autophagy.

Chromatin in *C. elegans* exhibits a conserved architecture, characterized by canonical histones (H2A, H2B, H3, H4), linker histone H1, and various non-histone chromatin proteins such as HPL-1/2, MET-2, and MES proteins (Von Diezmann & Rog 2021). Our immunoprecipitation data demonstrate selective interaction of T13F2.2 with core histones H2A and H3 and the linker histone H1, but not H2B or H4. This pattern points toward a role in heterochromatin compaction and chromatin boundary maintenance.

Histone modification analysis revealed increased H3K9Ac and decreased H3K27me3 levels following T13F2.2 depletion, consistent with a transition from repressive to active chromatin state. Supporting this, our electron microscopy studies illustrated dramatic nuclear decompaction and loss of peripheral heterochromatin. The observed nuclear envelope distortions and altered nucleolar integrity reflect compromised chromatin tethering and architectural integrity—hallmarks of aged or stressed nuclei.

These findings align with studies reporting that chromatin relaxation predisposes cells to genome instability and premature senescence, emphasizing the role of T13F2.2 in preserving nuclear integrity via epigenetic regulation. Our data underscore functional parallels between T13F2.2 and human PC4 (positive coactivator 4). Despite modest sequence identity, structural modeling, immunoreactivity with anti-PC4 antibody, and nuclear localization in heterochromatic foci establish a conserved molecular role. Human PC4 is implicated in transcription initiation, chromatin compaction, and autophagy inhibition (Sikder et al. 2019). The localization of T13F2.2 at the nuclear periphery, its interaction with histones, and the transcriptomic upregulation of chromatin remodeling and autophagy genes upon its silencing confirm these functions.

However, certain distinctions exist. For instance, while PC4 has been extensively studied in human systems, the cellular and developmental context in *C. elegans* is more transparent and tractable. Our results represent the first *in vivo* demonstration of a PC4-like protein influencing lifespan and stress resilience through nuclear architecture remodeling. This expands the scope of PC4-like proteins into invertebrate aging models and positions T13F2.2 as a functional ortholog with evolutionary divergence.

This study identifies T13F2.2 as a critical chromatin-associated protein that governs nuclear architecture, modulates the redox-autophagy axis, and ultimately determines organismal longevity in *C. elegans*. Its functional convergence with the human coactivator PC4 uncovers new dimensions in the chromatin-autophagy-aging interplay (Sikder 2019, Mustafi et al. 2022). By revealing how a single nuclear protein can orchestrate epigenetic remodeling, cellular stress responses, and lifespan, this work lays the foundation for further investigations into non-histone chromatin regulators as potential modulators of aging and age-related diseases and identifies a novel PC4 homologue in *C. elegans*, which plays a key role in organismal longevity.

## Materials and Methods

### *C. elegans* strains and culture

*C. elegans* were cultured on nematode growth media (NGM) and fed with the E. coli OP50 strain (uracil auxotroph) bacteria on lawns maintained at 22°C (Jadiya et al. 2016). To obtain a synchronized worm population, mature gravid worms were treated with 4% sodium hypochlorite. Strains used included the N2 strain (wild type), DA2123 ([lgg-1p::GFP::lgg-1 + rol-6(su1006)]), BC1292 (sIs10729 [rCes T12G3.1::GFP + pCeh361]), and J2586, cox-4(zu476[cox4::eGFP::3xFLAG]) (Jenzer et al. 2015). *C. elegans* strains were sourced from the Caenorhabditis Genetics Center (University of Minnesota).

### RNAi-Induced Gene Silencing

RNAi of the target gene was achieved using the feeding method as previously described (Jadiya et al. 2016). RNAi clone bacteria in the HT115 (*E. coli* strain) were grown in LB broth containing 50 μg/ml ampicillin. This culture was then incubated at 37°C for 6 hours, allowing the RNAi bacterial clones to reach the log phase for optimal RNAi efficiency. Individual RNAi bacterial clones were seeded on NGM plates with 5 mM isopropylthio-beta-D-galactoside (IPTG) for siRNA induction and 25 mg/ml carbenicillin as a selection agent. The plates were then incubated at 22°C overnight (12 to 14 hours) and used for subsequent experiments the following day.

### Life span

Life span experiments were conducted using a standard method and analyzed with the Kaplan-Meier survival curve (Dallaire et al. 2014). Briefly, 100 young adult worms were transferred to a new RNAi plate 48 hours post-embryo isolation. These worms were transferred to fresh plates until the last one perished. To maintain synchronization, the worm population’s data was analyzed using the Prism software, employing the survival curve graph.

### Estimation of Reactive oxygen species (ROS)

To gauge the effect of RNAi treatments on ROS levels in worms, the 2,7-dichlorodihydrofluorescein-diacetate (H2DCFA) assay was employed (Sharma et al. 2018). H2-DCFDA dye penetrates living cells, where intracellular esterase cleaves diacetate, converting it to H2-DCA. This H2-DCA then reacts with a host cell’s ROS, oxidizing into 2,7-DCF. The fluorescence of 2,7-DCF, indicating the relative concentration of ROS levels inside the cell, is observed with an excitation wavelength of 495 nm and an emission range of 512-527 nm. For our study, the N2 strain was utilized to examine the effects of the target gene’s RNAi on ROS levels. RNAi-treated worms, at the young adult stage, were harvested, washed three times with M9, and once with 1X PBS. Samples containing 100 worms each were pipetted into 96-well plates in triplicate. The volume was adjusted to 100 μl with 1X PBS. Absorbance readings were taken at the requisite wavelengths. An initial reading determined the buffer’s baseline absorbance. A subsequent reading was taken post the addition of 100 μl of 100 μM H2DCFA, with the final reading taken an hour later. ROS levels in individual worms were determined by subtracting the initial absorbance reading from the 0-hour reading post-dye addition, and then from the 1-hour reading. The resulting data was plotted, showcasing the relative fluorescence intensity per worm compared to controls.

### Transmission electron microscopy

TEM was executed as previously described, albeit with several adjustments (Kovács 2015; Kathuria et al. 2014). *C. elegans* worms, preserved in a phosphate buffer containing 2.5% glutaraldehyde and 4% paraformaldehyde, were incised at both ends to ensure swift penetration of the fixative, solutions, and resin infiltration, a process otherwise hindered by their dense, impermeable cuticle. A small volume of 0.1% ruthenium red was incorporated into the fixative and left overnight at 4°C. Worm segments were then encased in 2% low-melting agarose and post-fixed in 2% osmium tetroxide. Subsequent dehydration occurred in a progressively concentrated series of ethanol and was ultimately embedded and polymerized in low-viscosity spur resin. Worm pieces were oriented in flat embedding molds to ensure longitudinal sectioning. Ultrathin sections, ranging between 60 and 80 nm in thickness, were prepared using a Leica UC7 ultra-microtome. These sections were collected on 200 mesh copper grids, stained with lead citrate and uranyl acetate, and viewed under a Jeol JEM 1400 TEM at 80 kV. Data collection was facilitated by a Gatan 2k x 2K Orius CCD camera, post-optimal optical alignment.

### Total RNA Isolation and cDNA Synthesis

Age-synchronized worms were washed three times with M9 buffer, followed by two washes with diethylpyrocarbonate (DEPC; Sigma, Cat. No. D5758) treated water to eliminate adherent bacteria (Sarkar, Hameed, et al. 2022). Total RNA from all *C. elegans* strains was extracted using the RNAzol® RT method (Sigma, Cat. No. R4533) following the manufacturer’s guidelines. RNA concentrations were ascertained using the Nanodrop UV spectrophotometer (Thermo, Quawell, UV-Visible Spectrophotometer, Q5000). Adhering to the manufacturer’s instructions, 1000ng of total RNA was employed for cDNA synthesis utilizing the Verso cDNA Synthesis Kit (Thermo Fisher; Cat no: AB1453B). The cDNA concentration was subsequently quantified in ng/μl using a Nanodrop spectrophotometer.

### Quantification of mRNA

mRNA levels for all genes were determined through real-time quantitative PCR, with comparative analyses conducted using the N2 (wild type) strains (Sarkar, Shamsuzzama, et al. 2022). The previously described method was utilized for initial total RNA extraction. In brief, worms were washed three times with 0.2% DEPC water. The sample was then treated with QIAzol Lysis Reagent, and cDNA synthesis was achieved using the Verso cDNA Synthesis Kit (Thermo Scientific). For the real-time PCR reactions, 100 ng of cDNA from each sample was employed, and amplification was facilitated using the SYBR Green q-PCR Thermo kit. For mRNA amplification, a 96-well plate was utilized, and the amplification program was set as follows:

a. Pre-incubation: One cycle at 50°C for 2 minutes and then 95°C for 10 minutes.
b. Amplification: 40 cycles with a sequence of 95°C for 30 seconds, 55°C for 30 seconds, and 72°C for 30 seconds.
c. Melting curve analysis: 95°C for 5 seconds and then 65°C for 1 minute.
d. Cooling: 40°C for 5 minutes.

The amplification protocol was adjusted based on the specific kit used. Comparative quantification of the target gene employed Ct values, with relative quantification founded on the 2-ΔΔCt method.

### The following are the details of the primers used in this study

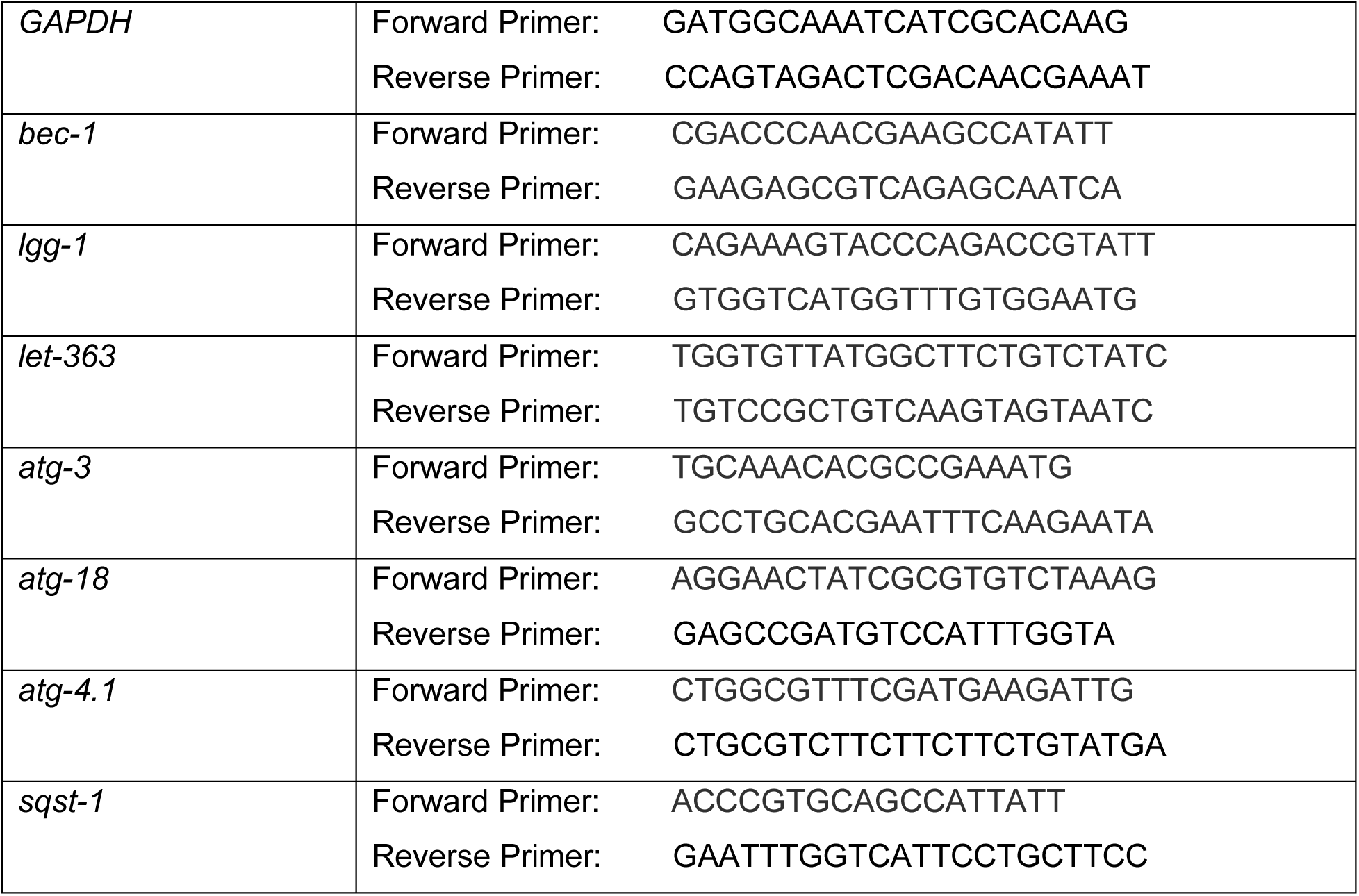
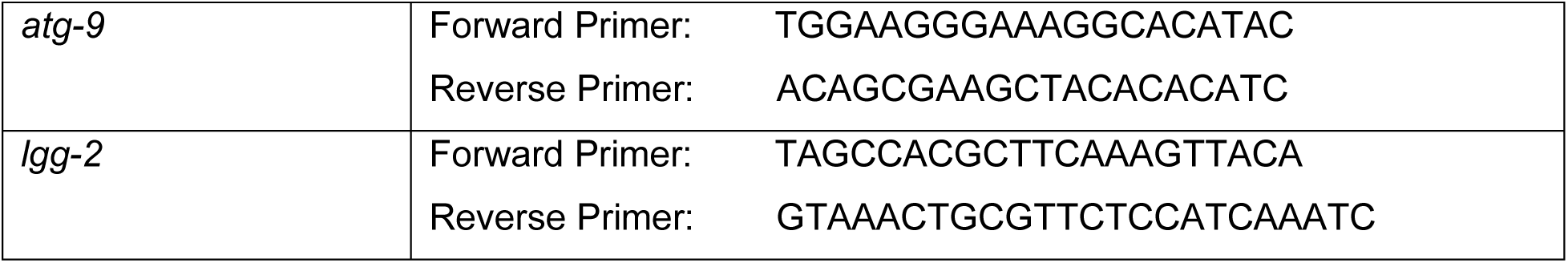

### Western Blotting

Worms were washed three times with M9 buffer and resuspended in worm lysis buffer containing 20mM potassium phosphate, 2mM EDTA, 1% Triton X-100, and 1x Protease K (Sarkar, Hameed, et al. 2022). Proteins were extracted from the worms either via sonication or by using the liquid N2 method in worm lysis buffer. Sonication followed a set protocol: 25 amplitude for 3 minutes with 15-second on-off pulse intervals using a Thermo Scientific sonicator. The sonicated lysate was centrifuged at 13,000 rpm for 20 minutes at 4°C, with the supernatant containing total worm protein. Protein concentration was determined using the Bradford reagent. For each group, 20-40 µg of protein was loaded onto 12-15% SDS-PAGE. The primary antibodies for western blotting were diluted at 1:1000, and the secondary antibodies were diluted at 1:1000 in 0.01% PBST. Chemiluminescence detection, western blot analysis, and densitometry measurements were carried out according to standard protocols using ImageJ software.

### Immunohistochemistry

We employed a modified standard IHC method (Crittenden & Kimble 2009). Nematodes from treatment plates were washed three times with M9 buffer to remove any adhering bacteria. The worms were then fixed in 4% paraformaldehyde prepared in PBS and left at 4°C overnight. Paraformaldehyde-fixed worms underwent freeze fracturing: they were placed between two microscope slides and exposed to liquid nitrogen for 10 minutes or kept at-45°C for two hours. The freeze-fractured worms were treated with a permeabilization solution (containing 5% fresh β-mercaptoethanol, 1% Triton X-100, and 125mM Tris, pH 7.4) for 48 hours at 37°C. After washing with 0.1% PBST, the worms were treated with 5% BSA for 3 hours as a blocking solution, using an end-to-end rotor in a 1.5 ml conical tube. Worms were then incubated overnight at 4°C with the respective primary antibody at a 1:200 dilution. The following day, after washing with 0.1% PBST, they were treated with secondary antibodies at a 1:500 dilution for 4 hours at 4°C. Finally, worms were centrifuged and mounted on microscope slides using 50% glycerol. Images were captured using a Leica confocal microscope at 60X and 100X magnification.

### Co-Immunoprecipitation

Worm extracts were prepared from wild-type N2 worms and subjected to immunoprecipitation using the mammalian PC4 antibody. For the immunoprecipitation experiment, 2 mg of total protein was used. The lysate was incubated overnight with 0.8 µg of human PC4 antibody at 4°C. The next day, Protein G-sepharose beads (Cat no. ab193259) were added to the lysate-antibody complex and incubated for 4-5 hours at 4°C. The mixture was then centrifuged at 2000 rpm for 3 minutes at 4°C. After washing the protein-antibody-bead complex three times with M9 buffer, the protein was eluted with 2X SDS dye. Immunoblotting was subsequently performed with the corresponding core-histone antibody to investigate histone interactions. Antibody details are as follows: H3K9ac: lab raised; H3K27me3 (ab6002); H3: lab raised; PC4: (ab154852); H1: (ab05-457); H2A: lab raised; H2B: lab raised and H4:lab raised.

### Whole Worm Transcriptome Profiling

Total RNA integrity was validated using an Agilent Bioanalyzer 2100, and only samples with distinct rRNA peaks were further processed. Libraries for RNA sequencing were prepared following the protocol provided with the KAPA stranded RNA-seq kit with RiboErase (KAPA Biosystems, Wilmington, MA, USA). The quality and quantity of the final libraries were assessed using the Agilent Bioanalyzer 2100 and the Life Technologies Qubit 3.0 Fluorometer, respectively. Sequencing was carried out in a paired-end 150 bp format on the Illumina HiSeq X platform (Illumina Inc., San Diego, CA, USA).

### RNA-sequencing analysis

Raw FASTQ reads were quality and adapter trimmed using Trim Galore! (https://github.com/FelixKrueger/TrimGalore) (version 0.6.7) with the paired option. Quality trimmed reads were aligned to the *C. elegans* reference genome (WBcel235/ce11) using HISAT2 (version 2.2.1) using very-sensitive no-soft clip options (Kim et al. 2019). Read pairs that did not map in a proper pair were removed, and reads that were secondary alignments were removed using the same tools (Danecek et al. 2021). Duplicate reads were marked, and optical duplicates were removed using Picard Mark Duplicates (https://broadinstitute.github.io/picard/). Gene counts were obtained against the ce11 Ref Seq gtf file using feature Counts (Liao et al. 2014). The DESeq2 package was used to identify differentially expressed genes (log2 fold change > 0.6 and padj < 0.1). The DAVID tool was used to determine significantly altered gene ontology and pathways (Huang et al. 2009). For displaying specific pathways, Worm Paths was used (Walker et al. 2021). Figures were made in R (version 4.2.2) using the ggplot2 (Wickham 2016) and heatmap packages (Kolde R (2018). *pheatmap: Pretty Heatmaps*. https://github.com/raivokolde/pheatmap) and assembled in Illustrator.

### Photomicrographs Recording and Analysis

Synchronized worms at the L4 stage underwent three wash cycles with M9 buffer and were immobilized using 100 mM sodium azide (Sigma, Cat No. 71289) (Sarkar, Shamsuzzama, et al. 2022). These immobilized specimens were positioned on glass slides, overlayed with a coverslip. Image capturing utilized a Carl Zeiss Axio Imager M3 microscope in tandem with ZEN2010 image acquisition software. Photomicrographs from each group (n=30, both 10x and 20x magnifications) were assessed using the Image J software from the National Institutes of Health, Bethesda, Maryland. A consistent analyzed area was maintained throughout all conditions. Mean fluorescence intensity values and statistical data were derived using GraphPad Prism 5.

## Data Analysis

Data representations were based on results from a minimum of two independent experiments, with the number of replicates as mentioned for each endpoint. The data were further presented as mean ± standard error of the mean. Statistical significance was determined via Student’s t-test, using GraphPad Prism 5, with a significance threshold set at p<0.05.

## Availability of data and materials

The data that support the findings of this study are available from the corresponding author upon reasonable request.

## Authors’ contributions

*CSIR-Central Drug Research Institute, Lucknow 226031, India; Academy of Scientific and Innovative Research (AcSIR), Ghaziabad 201002, India*

RH conducted experiments, analyzed data, and wrote the manuscript. KS did the structural analysis of T13F2.2. KM performed the transmission electron microscopy experiments and analyzed the data. ANazir conceived the study, provided infrastructure and reagents, analyzed the data and edited the manuscript.

*^2^Transcription and Disease Laboratory, Molecular Biology, and Genetics Unit, Jawaharlal Nehru Centre for Advanced Scientific Research, Bengaluru 560 064, India;*

Sweta Sikder, Aayushi Agrawal conducted some of the experiments. TKK conceived the study, provided infrastructure and reagents, analyzed the data, and edited the manuscript.

*^3^Theomics International Private Limited Bangalore, India*

Madavan Vasudevan performed the total RNA Sequencing.

*^4^Cancer Research Division, Rajiv Gandhi Centre for Biotechnology (BRIC-RGCB), Thiruvananthapuram, Kerala, 695014, India*.

Parijat Senapati analyzed the RNA Sequencing Data.

## Acknowledgements

The SAIF facility of CSIR-CDRI was used for carrying out some experiments and is gratefully acknowledged. TKK acknowledges the central facility of MBGU, JNCASR, Bengaluru 560064, India. CSIR-CDRI Communication Number: 177/2025/AN&TKK.

## Supplementary Information

**Fig-SI-I, (a-f):**
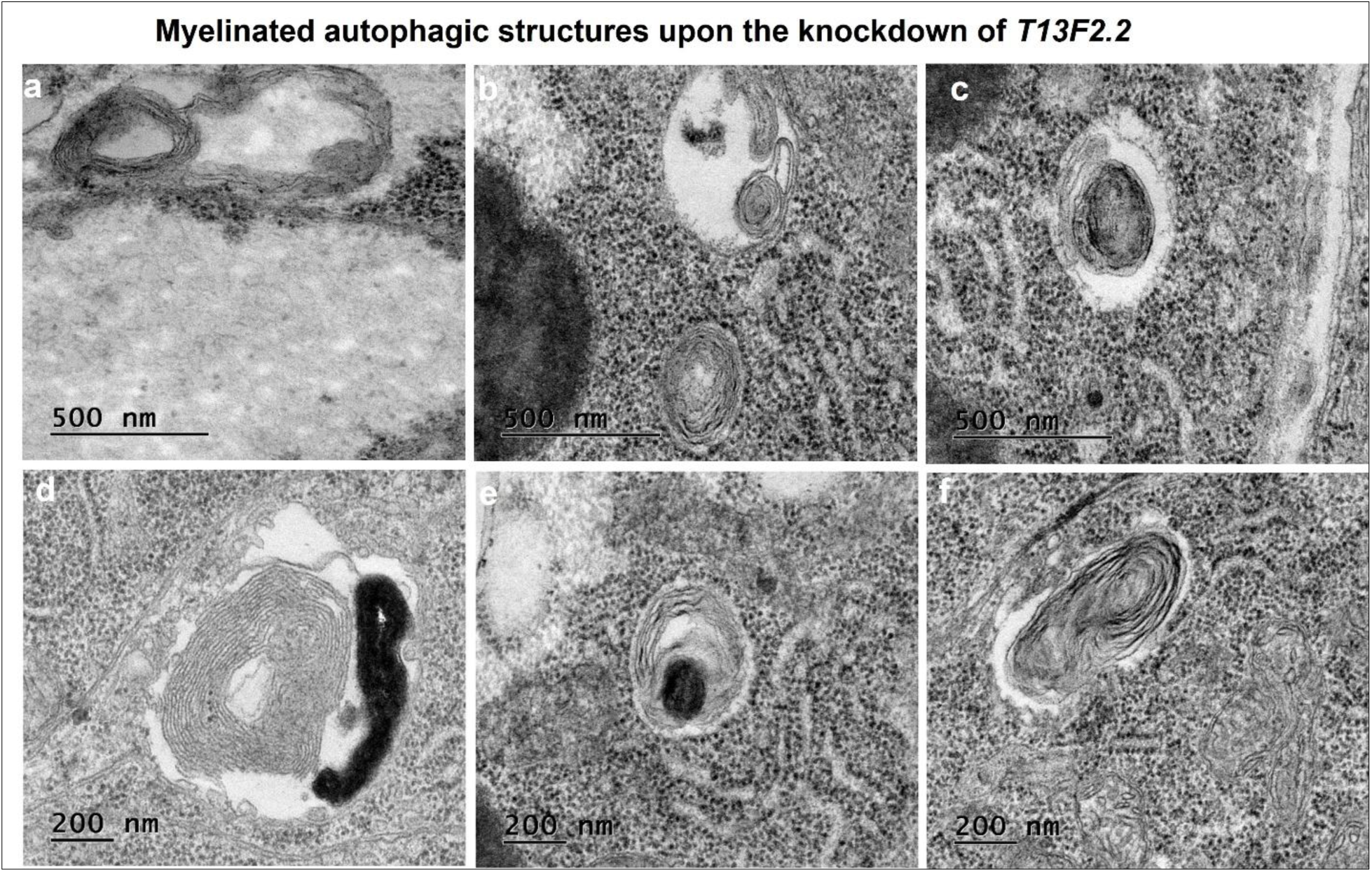
TEM photomicrographs of N2 worms (wild type) upon the knockdown *T13F2.2*, show multiple myelinated autophagic structures.

**Fig-SI-II, (a-d):**
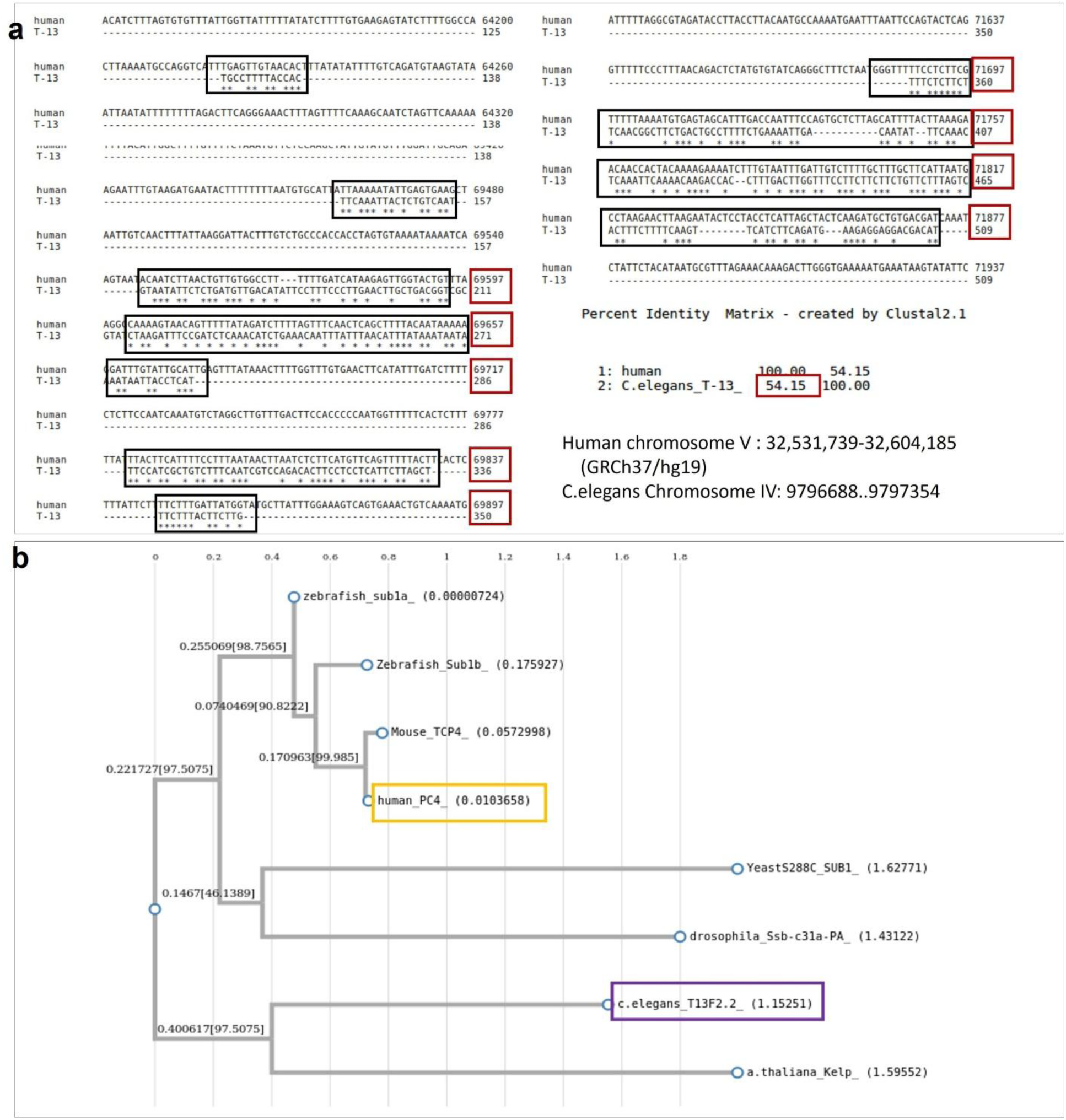
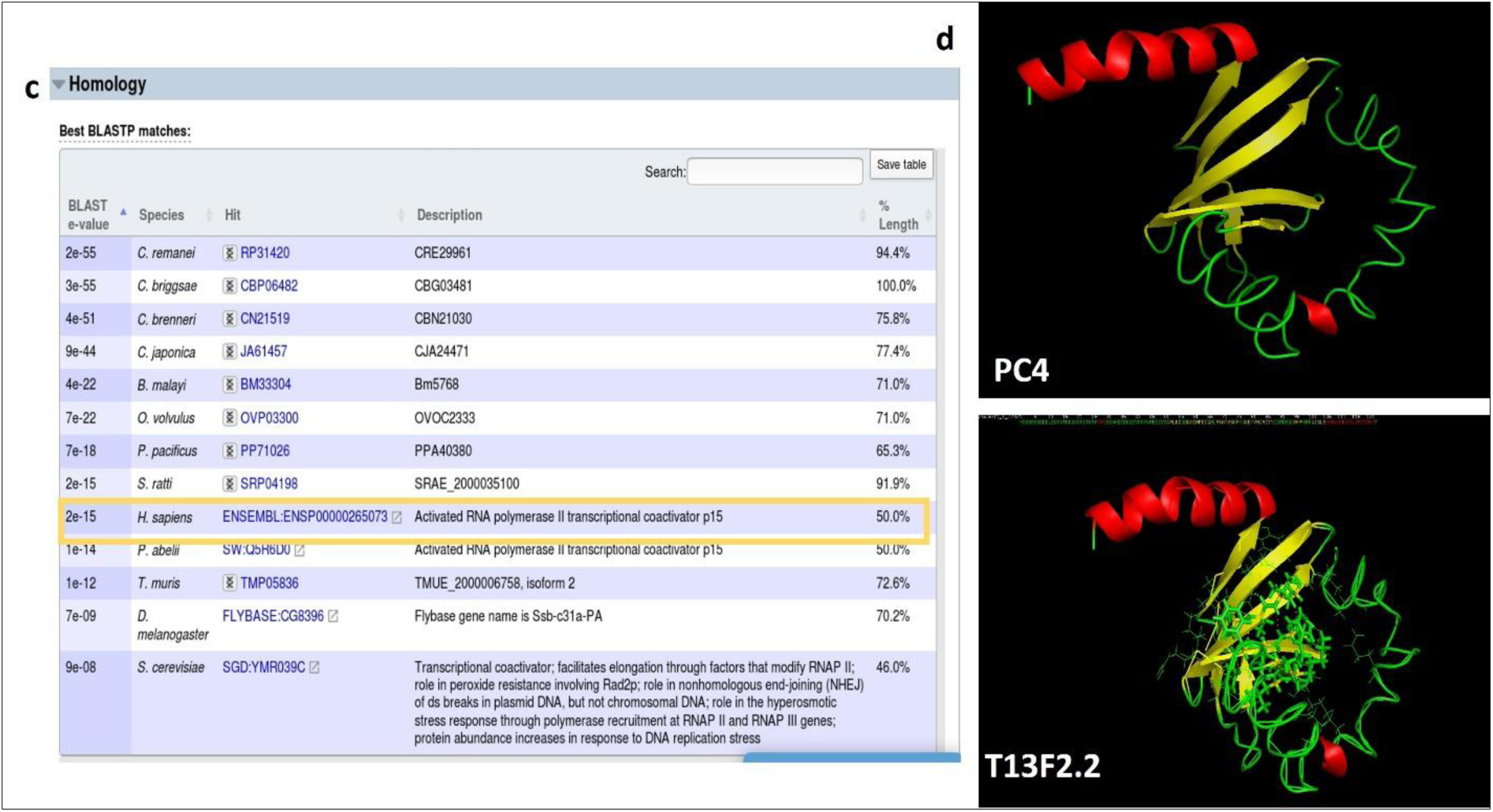
*in silico* structure prediction of *T13F2.2*, shows human and *C. elegans* protein conserved C-terminal domain with C-Score: Confidence score [-5 2]: the predicted model is of moderate confidence, TM-Score: Sequence similarity with the template is 39%.

**(Fig-SI-IV):**
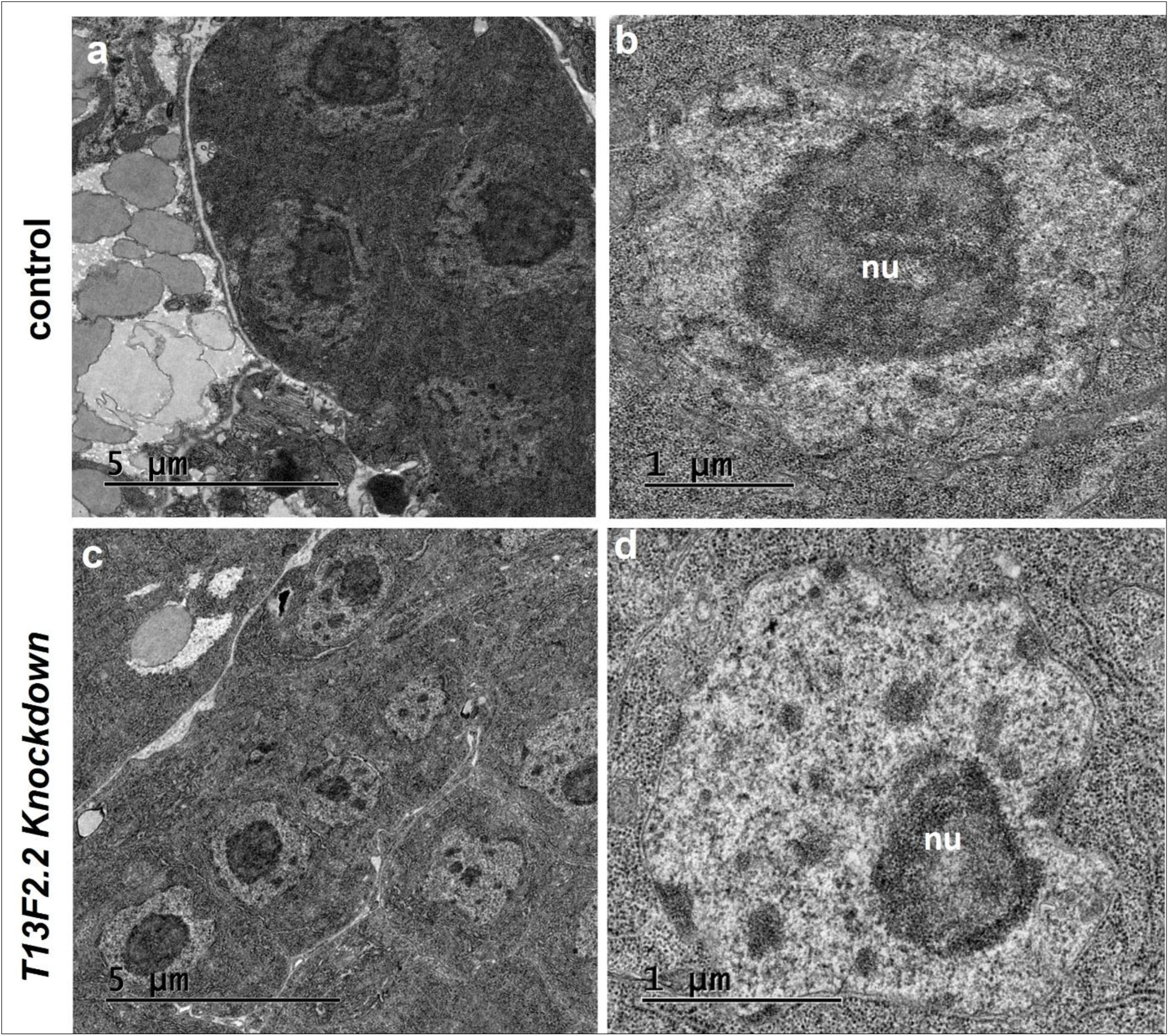
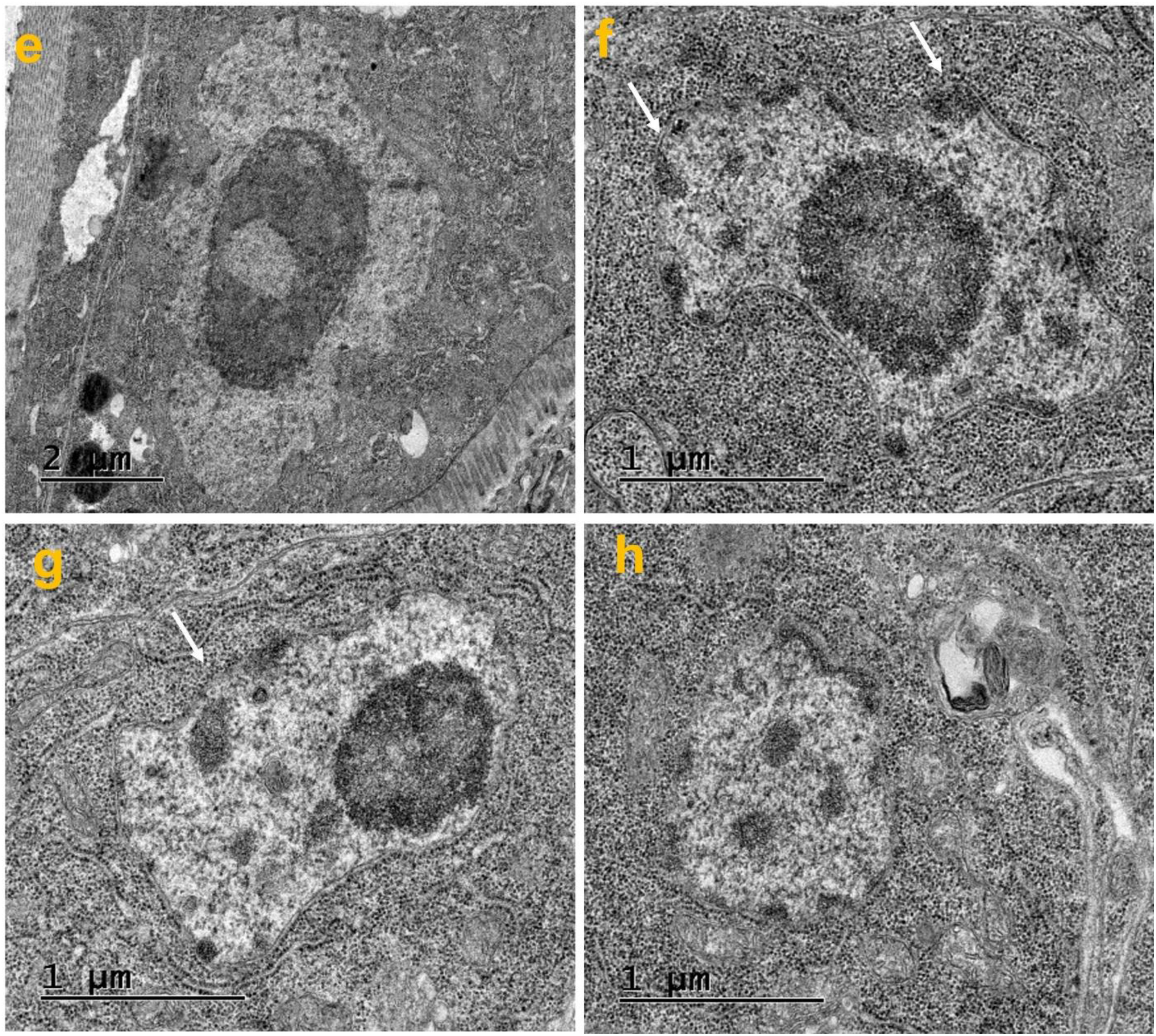
(a-b) Control worms show normal spherical nucleus with well distributed heterochromatin, nucleolus (Nu), and well-preserved nuclear envelope. **(C-J)** Transmission electron micrographs reveal fine structures of nuclei from control and knockdown worms; *T13F2.2* knockdown nucleus is more lobulated and deformed with disruption of the nuclear envelope (arrows) along with loss of heterochromatin from nuclear periphery.

